# Integrin-dependent migratory switches regulate the translocation of *Toxoplasma*-infected dendritic cells across brain endothelial monolayers

**DOI:** 10.1101/2021.04.21.440681

**Authors:** Emily C. Ross, Arne L. ten Hoeve, Antonio Barragan

**Affiliations:** Department of Molecular Biosciences, The Wenner-Gren Institute, Stockholm University, Stockholm, Sweden

**Keywords:** leukocyte, blood-brain barrier, Apicomplexa, immune cell, cell migration, cell adhesion molecule

## Abstract

Multiple cellular processes, such as immune responses and cancer cell metastasis, crucially depend on interconvertible migration modes. However, knowledge is scarce on how infectious agents impact the processes of cell adhesion and migration at restrictive biological barriers. In extracellular matrix, dendritic cells (DCs) infected by the obligate intracellular protozoan *Toxoplasma gondii* undergo mesenchymal-to-amoeboid transition (MAT) for rapid integrin-independent migration. Here, in a cellular model of the blood-brain barrier, we report that parasitised DCs adhere to polarised endothelium and shift to integrin-dependent motility, accompanied by elevated transendothelial migration (TEM). Upon contact with endothelium, parasitised DCs dramatically reduced velocities and adhered under both static and shear stress conditions, thereby obliterating the infection-induced amoeboid motility displayed in collagen matrix. The motility of adherent parasitised DCs on endothelial monolayers was restored by blockade of β1 and β2 integrins or ICAM-1, which conversely reduced motility on collagen-coated surfaces. Moreover, parasitised DCs exhibited enhanced translocation across highly polarised primary murine brain endothelial cell monolayers. Blockade of β1, β2 integrins, ICAM-1 and PECAM-1 reduced TEM frequencies. Finally, gene silencing of the pan-integrin-cytoskeleton linker talin (*Tln1*) or of β1 integrin (*Itgb1*) in primary DCs resulted in increased motility on endothelium and decreased TEM. Adding to the paradigms of leukocyte diapedesis, the findings provide novel insights in how an intracellular pathogen impacts the migratory plasticity of leukocytes in response to the cellular environment, to promote infection-related dissemination.

## Introduction

Dendritic cells (DCs) are central to mount immune responses and to combat infections [1]. Their effector functions are crucially linked to a remarkable migratory plasticity, for example the ability to transition from mesenchymal to amoeboid (MAT) migration modes [2]. This requires major changes in the actin cytoskeleton, and enables high-speed locomotion through extracellular matrix (ECM), when travelling from the periphery to secondary lymphoid organs, or to reach biological barriers, on their way from the blood into tissues [3,4]. Amoeboid migration is especially suited for rapid locomotion of leukocytes in cellular networks and tissues [5]. In contrast to integrin-dependent mesenchymal migration on 2-dimensional (D) surfaces, amoeboid migration in 3D confinements occurs independently of adhesion to specific substrates or ECM degradation and relies primarily on the protrusive flow of actin filaments for efficient migration through interstitial tissues [6-8]. However, upon reaching biological barriers, DCs in the circulation also need to adhere to the vascular endothelium.

The blood-brain barrier (BBB) regulates traffic of leukocytes and protects the brain from pathogens, toxins, cell damage and inflammation [9]. Specialised endothelial cells, held together by tight junctions (TJs), form a highly restrictive barrier that tightly regulates homeostasis in the central nervous system (CNS) [10]. The cerebral endothelium is also in constant crosstalk with astrocytes, microglia, neurons, pericytes and circulating immune cells [11], that all contribute to protect the CNS from external insults.

Upon inflammation, recruitment of leukocytes, such as DCs, and their interaction with the vascular endothelium is a critical step in the immune response [12]. Endothelial cells attract and direct leukocytes to enter underlying tissue, by exposing a variety of cell adhesion molecules (CAMs), such as ICAM-1 and PECAM-1, at their surface that capture travelling immune cells in the bloodstream [13]. Leukocytes on the other hand, express various integrins on their surface, which include β1 and β2 integrins that bind to CAMs on the endothelial cell surface [14]. Talin-1 connects integrins to actin and contributes to the regulation of integrin adhesion complexes [15]. Jointly, these interactions mediate firm adhesion and spreading of leukocytes on the endothelium, which leads to leukocyte crawling and eventually transmigration through the endothelial barrier to reach sites of inflammation [16].

Understanding the biology of leukocyte migration can be aided through the study of host-pathogen interactions, owing to the microbial ability to hijack host cell functions. *Toxoplasma gondii* is a model parasite [17] and obligate intracellular pathogen that infects warm-blooded vertebrates. It is estimated that one third of the global human population encounters *T. gondii* during a life-time [18]. *T. gondii* has a remarkable ability to cross non-permissive cellular barriers to reach immunoprivileged sites, such as the CNS [19]. Primary infection is usually subclinical, and once the parasite reaches the CNS, it can chronically persist inside cysts. In immunocompromised individuals or in the developing foetus, reactivation of latent infection or acute disseminated infection, respectively, can be life-threatening and cause encephalitis [20].

Mononuclear phagocytes, such as DCs, can act as ‘Trojan horses’ for *T. gondii* dissemination in mice [21-23]. When actively invaded by *T. gondii*, DCs undergo cytoskeletal changes with redistribution of integrins, reminiscent of MAT in ECM [24-26]. This migratory activation -termed hypermotility- [27] requires live intracellular parasites [25], does not rely on chemotaxis [28] and is instead mediated by non-canonical GABAergic signalling and MAP kinase activation [23,29-32]. However, it has remained unknown how highly-migratory parasitised DCs interact with polarised endothelium.

In the present study, we addressed the interaction of parasitised DCs with polarised primary brain endothelial cell monolayers, and identified a migratory shift associated to transendothelial migration (TEM), with a central role for integrins and CAMs.

## Materials and Methods

### Parasite culture and cell lines

The GFP or RFP-expressing PTG/ME49 *T. gondii* strains (type II) [33,34] were maintained by serial 48 h passaging in human foreskin fibroblasts (HFFs; CRL-2088, American Type Culture Collection). HFFs were cultured in Dulbecco’s modified Eagle’s medium (DMEM, Thermofisher scientific) with 10% heat inactivated fetal bovine serum (FBS, Sigma), gentamicin (20 μg/ml, Gibco), L-glutamine (2 mM, Gibco) and HEPES (10 mM, Gibco), referred to as D10. bEnd.3 cells (CRL-2299, American Type Culture Collection) were cultured in D10. All cell cultures and parasites were grown in a humidified atmosphere containing 5% CO2 at 37°C.

### Primary dendritic cells (DCs) and macrophages

Murine bone marrow-derived DCs were generated as previously described [28]. Briefly, cells from bone marrow of 6 to10-week old male or female C57BL/6NCrl mice (Charles River) were cultivated in RPMI 1640 with 10% fetal bovine serum (FBS), gentamicin (20 μg/ml), glutamine (2mM) and HEPES (0.01 M), referred to as complete medium (CM), and supplemented with recombinant mouse GM-CSF (20 ng/ml, Peprotech). Loosely adherent cells (DCs) were harvested after 6 or 8 days. To generate macrophages, cells from bone marrow were cultivated in CM supplemented with M-CSF (20 ng/ml, ImmunoTools). Loosely adherent cells were discarded and adherent cells were harvested after 6 or 8 days.

### Primary murine brain endothelial cells (MBECs)

Six to 8-week old C57BL/6NCrl mice, housed under specific pathogen-free conditions at Stockholm University, were euthanised and brains were extracted. The brain tissue was freed from the cerebellum, hypothalamus, olfactory bulb and kept in ice-cold PBS. Preparation of cells was performed as described [35], with modifications [36]. The tissue was digested in collagenase IV (1 mg/ml, Gibco) for 1 h at 37° C, homogenised and washed with wash buffer (PBS, 0.5% FBS and 2mM EDTA). The tissue suspension was mixed with density gradient medium. Percoll (Easycoll, Biochrom) in PBS was added to the tissue suspension at a final concentration of 30% and centrifuged for 30 min at 500 g. After removal of the myelin on the top layer, the cell pellet was washed with wash buffer. A second digestion was carried out by incubating the cell pellet with collagenase IV for 30 min at 37° C. Positive selection of CD31-expressing cells was performed by magnetic activated cell sorting (MACS, Myltenyi Biotec) and CD31 MicroBeads (Myltenyi Biotec), according to the manufacturer’s instructions. CD31^+^-enriched cell suspensions were finally plated onto transwell filters (8 µm pore size; BD Biosciences) at cell yield from 1 brain per 4 transwell inserts pre-coated with 0.1% gelatin (Gibco), or in 12-well culture plates and monitored for confluence and polarisation. Cells were cultured in EBM-2/EGM-2 medium (Lonza) with 12% FBS, glutamine (200 mM, Gibco) and growth supplements (Bulletkit™, Lonza) minus the vascular endothelial growth factor (mVEGF) supplement. EBM-2/EGM-2 medium was supplemented with puromycin (1 mg/ml, Gibco) for 3 d, after which medium was changed every 2 d in absence of puromycin. The cellular monolayers had high and stable expression of PECAM1/CD31 (>99%), TJ markers ZO-1, occludin and claudin-5, with low expression of astrocyte marker GFAP (<1%), as previously characterised [36].

### Polarisation parameters: permeability assay and transendothelial electrical resistance (TEER)

bEnd.3 cells were cultured to 80% confluence then seeded onto transwells (8 µm pore size; BD Biosciences) and grown for 5 d until they reached polarisation, as defined below. MBECs were seeded directly after isolation and reached a polarisation plateau after ∼ 11-12 d, which was maintained beyond 14 d, as defined below. Experiments were performed on d 13-14. For evaluation of cell monolayer permeability following treatments or transmigration, FITC-dextran (3 kDa; Life tech) was added to the upper compartment of the transwell at a concentration of 12.5 μg/ml for 90 min. Medium was collected from the lower compartment, and fluorescence was measured in a fluorometer (EnSpire Multimode Plate Reader, Perkin Elmer) at 485 nm excitation 520 nm emission.

MBECs and bEnd.3 cells were cultured to form polarised monolayers defined by a TEER ≥ 250 Ω • cm^2^, as measured using an Ohmmeter (Millipore, Bedford, MA) and correcting measurements with the formula: Unit Area Resistance (TEER) = Resistance (Ω) • Effective Membrane Area (cm^2^). TEER was measured before and after transmigration or treatments. Values are shown as percentage (%) of TEER related to TEER prior to treatment or transmigration.

### Reagents

Blocking antibodies, isotype controls and inhibitor of VLA-4 were used at 1 μg/ml: LEAF™ purified anti-CD29 (clone HMβ1-1, 102209, BioLegend), LEAF™ purified anti-CD31 (clone MEC13.3, 102511, BioLegend), LEAF™ purified anti-CD18 (clone M18/2, 101409, BioLegend), LEAF™ Purified Rat IgG2a, κ Isotype Ctrl (clone RTK2758, 400515, BioLegend), Ultra-LEAF™ Purified Armenian Hamster IgG Isotype Ctrl (clone HTK888, 400969, BioLegend), BIO 5192 (R&D systems), anti-CD54 (ICAM-1; clone YN1/1.7.4; eBioscience).

### Immunostainings

*T. gondii*-challenged DCs were plated on coverslips coated with bovine collagen I (1 mg/ml, Life Technologies). After fixation (4% PFA, Sigma-Aldrich), cells were permeabilised (0.5% Triton X-100, Sigma-Aldrich) and stained with phalloidin Alexa Fluor 595 (Invitrogen). Micrographs were generated using a 63x objective (DMi8, Leica Microsystems). Primary MBECs were seeded on coverslips pre-coated with 0.1 % gelatin (BioRad). Fixation and permeabilisation steps were carried out as for DCs, followed by blocking (5% FBS in PBS for 2 h). Cells were then incubated with primary antibodies to ZO-1 (Thermofisher) and Occludin (Thermofisher) ON at 4° C at 1:500. Cells were then stained with Alexa Fluor 594-conjugated secondary antibodies (Invitrogen) and DAPI for 2 h, mounted and imaged by confocal microscopy (LSM 800, Zeiss).

### Transmigration assays

Day 6-8, DCs or macrophages were challenged with freshly egressed tachyzoites (ME49/PTG-GFP or ME49-RFP, MOI 2, 4 h), resulting in 60-70% infection frequency and ∼1,2 tachyzoites / infected cell [22]. Cells were then transferred to transwell filters (8 μm pore size; BD Biosciences), with pre-cultured polarised monolayers of bEnd.3 cells in CM or MBECs in EBM-2/EGM-2. After 16 h, transmigrated DCs were put on ice for 1 h and macrophages trypsinised (TrypLE™ Express; Gibco) for 10 min to disassociate adherent cells. Cells were then resuspended and counted in a Bürker chamber.

### Motility assays

Motility assays were performed as previously described [25]. Briefly, DCs were cultured with CM ± freshly egressed *T. gondii* tachyzoites (ME49/PTG-GFP or ME49-RFP, MOI 3, 4 h, resulting in 70-80% infection frequency) and with soluble reagents as indicated. DCs were then added to 96-well plates pre-cultured with bEnd.3 cells or embedded in bovine collagen I (1 mg/ml, Life Technologies). Live cell imaging was performed for 1 h, 1 frame/ min, at 10x magnification (Z1 Observer with Zen 2 Blue v. 4.0.3, Zeiss). Time-lapse images were consolidated into stacks and motility data was obtained from 30 cells/condition (Manual Tracking, ImageJ) yielding mean velocities (Chemotaxis and migration tool, v. 2.0). Infected cells were defined by GFP^+^ or RFP^+^ cells, as indicated.

### Flow condition assays

DCs were cultured as stated in motility assay and then added to fluidic channels (μ-Slide VI^0.4^; Ibidi) with confluent bEnd.3 cell monolayers or pre-coated with collagen I (1 mg/ml, Life Technologies), and allowed to adhere for 10 min. Phase-contrast and fluorescence images were first captured in static condition. Fluidic shear stress was then applied by flowing CM at 0.2 dyn/cm^2^ through the channels. Live cell imaging was immediately initiated and images acquired every 5 seconds for up to 10 min, at 10x magnification. Fluidic shear stress was then increased to 1 dyn/cm^2^ and live cell imaging immediately initiated. Time-lapse images were consolidated into stacks and motility and path-length data obtained from 20 cells (Manual Tracking, ImageJ) yielding mean velocities and pathlengths (Chemotaxis and migration tool, v .2.0). The percentage of remaining adherent cells was calculated by dividing the number of cells per frame following shear stress, by the number of cells in the same frame in static condition. Representative track plots were created in Python 3.6.9 (https://www.python.org/) with pandas 1.1.5 and matplotlib 3.1.3.

### Flow cytometry

Bone marrow-derived DCs were cultured in CM ± freshly egressed *T. gondii* tachyzoites (ME49/PTG-GFP, MOI 1) or LPS (100 ng/ml, serotype 011:B4, Sigma-Aldrich) for 4 or 24 h. Cells were stained on ice in FACS buffer (1% FBS and 1 mM EDTA in PBS) with Live/Dead Violet (L34955, Life technologies), anti-CD11c (clone N418, 25-0144-82, eBioscience) and anti-CD18 (clone M18/2, 101407, BioLegend), -CD29 (clone HMβ1-1, 102213, BioLegend) -CD54 (clone YN1/1.7.4, 116119, BioLegend) or isotype control antibodies (clone R35-95, 553930, BD Pharmingen; clone HTK888, 400924, BioLegend; clone RTK4530, 400611, BioLegend), fixed with 2% PFA and analysed on a BD LSRFortessa flow cytometer (BD Biosciences) with FlowJo software (v. 10, FlowJo LLC). Prior to staining, cells were blocked in FACS buffer supplemented with anti-CD16/CD32 antibody (Fc Block, BD Pharmingen).

### Lentiviral vector production and *in vitro* transduction

Self-complementary hairpin DNA oligos targeting *Itgb1* (shITGB1, TRCN0000066645, Genscript) or *Tln1* (shTln1, TRCN0000294832, Genscript) mRNA were on self-inactivating lentiviral vectors (pLL3.7) with eGFP reporter expression (**Table S1**). Transfer plasmid (shRNA targeting ITGB1, Tln1 or Luc) was co-transfected with psPAX2 (12260, Addgene) packaging vector and pCMV-VSVg (8454, Addgene) envelope vector into Lenti-X 293T cells (Clontech) using Lipofectamine 2000 (Invitrogen). The resulting supernatant was harvested 24 h and 48 h post-transfection. Supernatants were centrifuged to eliminate cell debris and filtered through 0.45-mm cellulose acetate filters. DCs (3 days post-bone marrow extraction) were transduced by adding lentiviral supernatants in the presence of DEAE dextran (8 µg/ml; Sigma-Aldrich) to cells for 4 h. After 5-7 days, transduction efficiency was examined for eGFP expression by epifluorescence microscopy (Z1 Observer with Zen 2 Blue v. 4.0.3, Zeiss) followed by expression analysis by qPCR for knockdown of targeted mRNA.

### Polymerase chain reaction (PCR)

Total RNA was extracted using TRIzol reagent (Sigma-Aldrich). First-strand cDNA was synthesised with Superscript IV Reverse Transcriptase (Invitrogen). Real-time quantitative polymerase chain reaction (qPCR) was performed using SYBR green PCR master mix (Kapa biosystems), forward and reverse primers (200 nM), and cDNA (100 ng) with a QuantStudio™ 5 real-time PCR system (Thermofisher). Glyceraldehyde 3-phopshate dehydrogenase (GAPDH) was used as a house-keeping gene to generate ΔCt values; 2^-ΔCt^ values were used to calculate relative knock-down efficiency. All primers (Invitrogen) were designed using the Get-prime or Primer-BLAST software (**Table S2**).

### Statistical analyses

All statistics were performed with Prism software (v. 8, GraphPad). In all statistical tests, values of *P* ≥ 0.05 were defined as non-significant and *P* < 0.05 were defined as significant.

## Results

### *T. gondii*-infected DCs transmigrate across polarised monolayers of primary mouse brain endothelial cells (MBECs) and bEnd.3 cells

To assess if *T. gondii*-infected DCs can cross polarising and TJ forming brain endothelial cell monolayers, primary MBECs from murine brains or bEnd.3 cells were grown to confluency and polarisation in a transwell system (**Fig. 1a, Supplementary Fig. S1**). Upon exposure to *T. gondii*-challenged DCs or unchallenged DCs, MBEC monolayers maintained low permeability to the low molecular weight tracer FITC-dextran (3kDa) (**Fig. 1b**) and high transendothelial electrical resistance (TEER) (**Fig. 1c**), consistent with characteristics of cerebral endothelium [36]. Importantly, *T. gondii*-challenged DCs transmigrated across polarised MBEC and bEnd.3 monolayers, at significantly higher frequencies compared with unchallenged DCs (**Fig. 1d, g**). In contrast, *T. gondii*-challenged macrophages transmigrated at significantly lower frequencies, with non-significant differences compared with unchallenged macrophages (**Fig. 1e, f, g**). We conclude that *T. gondii*-challenged DCs transmigrate across polarised endothelial monolayers, with undetectable perturbation of cellular barrier integrity.

**Figure 1.**
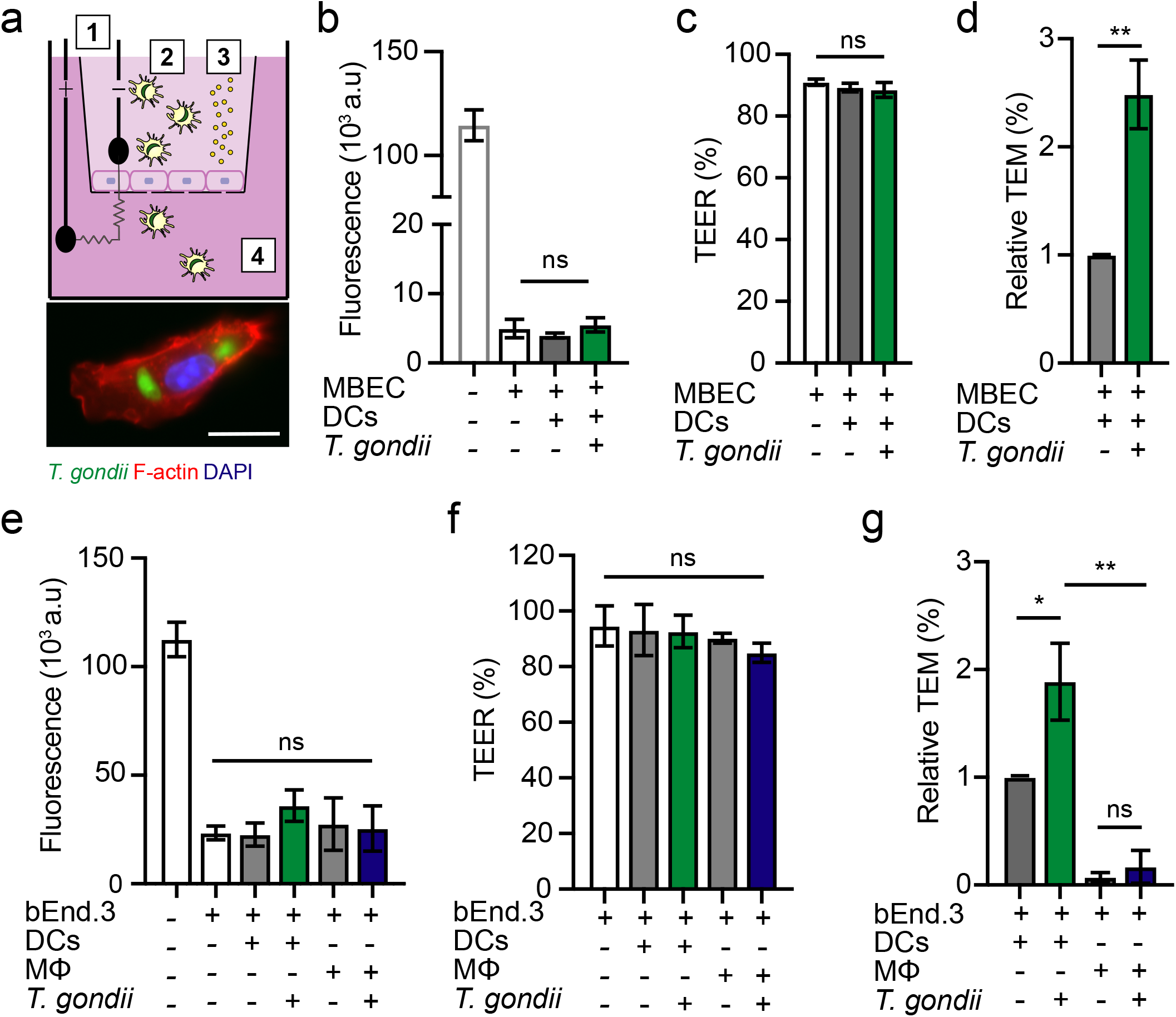
Transmigration of *T. gondii*-challenged DCs across polarised MBEC and bEnd.3 monolayers. **(a)** Top cartoon shows experimental setup in a 2**-**chamber transwell system and assessment of endothelial cell polarisation and permeability, as indicated in Materials and Methods. (1) Measurement of transcellular electrical resistance (TEER), (2) *T. gondii*-challenged or unchallenged cells were added to upper chamber, (3) The permeability tracer, FITC**-**dextran (3 kDa), added to upper chamber and measured in lower chamber, (4) Transmigrated cells were collected and counted. Bottom micrograph represents a DC infected with GFP-expressing *T. gondii* (green) and stained for F-actin (red) and nuclei (blue). Scale bar = 10 μm. **(b)** Permeability of MBEC monolayers to FITC-dextran (3 kDa) following DC transmigration. Data is shown as arbitrary fluorescence units (a.u). **(c)** TEER under same conditions as in (b). For each condition, TEER values (Ω•cm2) relative to TEER values at the initiation of the assay (100%) are indicated, as described in Materials and Methods. **(d)** Relative transendothelial migration (TEM) frequency of DCs across MBEC monolayers shown as percentage (%) of DCs added in the upper well, normalised to unchallenged DCs (set to 1.0). **(e)** Permeability of bEnd.3 cell monolayers to FITC-dextran (3 kDa) following transmigration of unchallenged or *T. gondii*-challenged dendritic cells (DCs) or macrophages (MF). **(f)** TEER values relative to TEER values at initiation of the assay (100%). **(g)** Relative transendothelial migration (TEM) frequency of DCs or MF across bEnd.3 monolayers shown as percentage (%) of DCs or MF added in the upper well, normalised to unchallenged DCs (set to 1.0). Bar graphs represent the mean ± s.e.m of 3-4 independent experiments (n = 3-4). \**P* < 0.05, \*\**P* < 0.01, ns: non-significant, by one-way ANOVA with Dunnett’s post-hoc test (b, c, e, f), Students’ *t*-test (d, g),

### Absence of hypermotility by *T. gondii*-infected DCs on endothelial cell monolayers

*T. gondii*-infected DCs are hypermotile in 2D and 3D confinements with collagen [23,30,26]. To determine their migratory behavior on endothelium, we compared the motility of DCs on bEnd.3 endothelial cell monolayers and on collagen-coated surfaces (**Fig. 2a**). In line with previous studies, *T. gondii*-infected DCs exhibited hypermotility on collagen, with elevations in migrated distances and velocity (**Fig. 2b, c**). In stark contrast, we found that *T. gondii*-infected DCs exhibited non-significant differences in velocity and migrated distances on bEnd.3 cell monolayers, compared with unchallenged DCs (**Fig. 2b, c**). The unexpected absence of hypermotility by *T. gondii*-infected DCs on endothelial cell monolayers motivated a further exploration of the cell-cell interactions in play.

**Figure 2.**
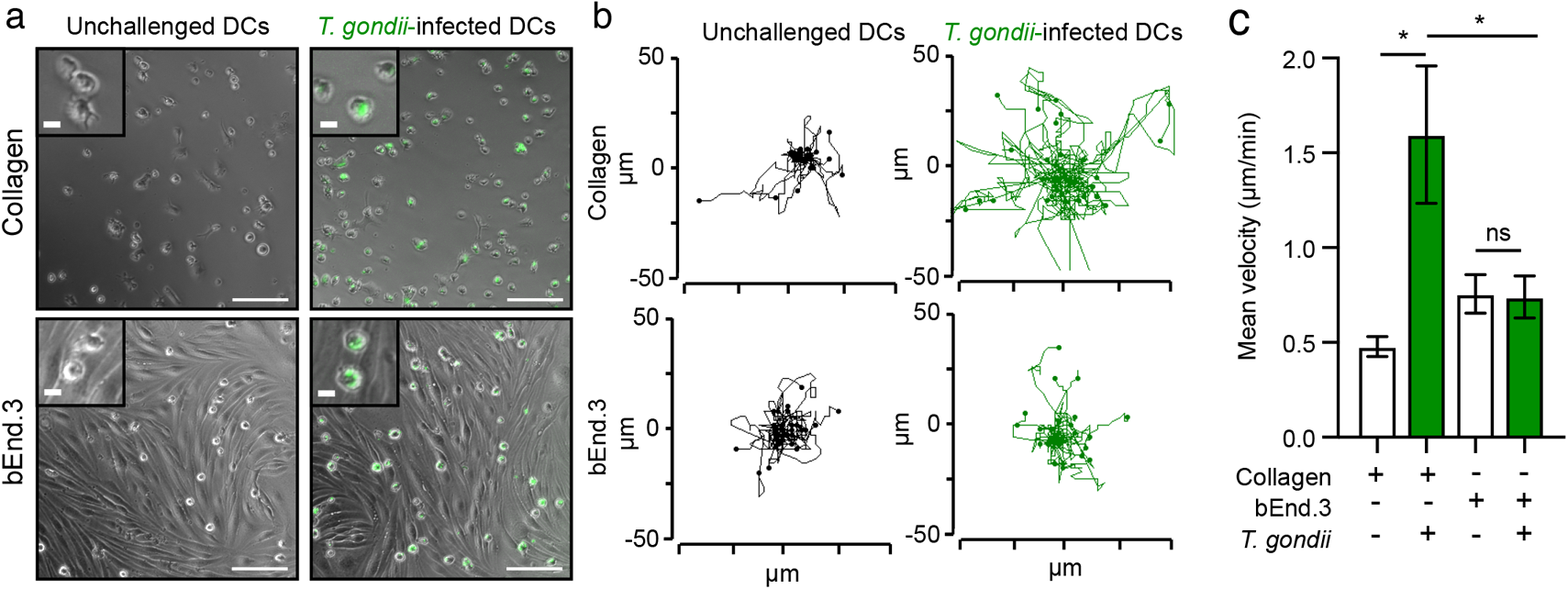
Motility analyses of *T. gondii*-infected DCs on collagen and endothelium. **(a)** Representative micrographs of unchallenged DCs and GFP-expressing *T. gondii*-challenged DCs on collagen (top row) and bEnd.3 cell monolayers (bottom row). Scale bar = 100 μm, inset = 10 μm. **(b)** Representative motility plots of DCs as described and represented in (a). **(c)** Mean velocity of unchallenged and *T. gondii*-infected DCs. Data represent the mean ± s.e.m from 3 independent experiments (n = 3). \**P* < 0.05, ns; non-significant by one-way ANOVA, Sidak’s post-hoc test.

### Integrins and CAMs are implicated in the motility of *T. gondii*-challenged DCs on endothelial cell monolayers

CAMs and integrins play central roles in adhesion, crawling and extravasation of leukocytes across endothelium [16]. We previously reported that integrin-blocking monoclonal antibodies (mAbs) non-significantly impacted the migration of parasitised DCs in a 3D collagen matrix [26]. To address the putative implication of integrins and CAMs on the motility of parasitised DCs on endothelium, we first assessed the expression of β1/β2 integrins and ICAM-1 upon *T. gondii* challenge by flow cytometry (Supplementary Fig. S2a, b). In parasitised DCs, the expression of β1/β2 integrins was maintained, with small elevations or non-significant differences compared to unchallenged DCs (Fig. 3a, b) or bystander DCs (Supplementary Fig. S2c, d) and with, overall, a relatively inferior response compared to stimulation with LPS (Fig. 3a, b; Supplementary Fig. S2c, d). Interestingly, the expression of ICAM-1 was significantly elevated by 24 h in *T. gondii*-infected DCs (Fig. 3c) but not in by-stander DCs (Supplementary Fig. S2d) Next, analyses of cell velocity were performed in presence of blocking mAbs. As expected, blockade of β1 and β2 integrins reduced the motility of parasitised DCs on collagen-coated surfaces (2D), with a non-significant impact by CAM blockade and antibody isotype controls (**Fig. 3d**). In sharp contrast, on bEnd.3 endothelial monolayers, blockade of ICAM-1 and β1 and β2 integrins conversely elevated the motility of *T. gondii*-challenged DCs (**Fig. 3e**). Blockade of PECAM-1, α4β1 integrin and isotype controls non-significantly impacted DC motility (**Fig. 3e**). Thus, ICAM-1- and β1- and β2-integrin blockades had opposite effects on DC velocities on collagen and endothelial cell monolayers (**Fig. 3f**). These dramatic and seemingly contraposed motility effects motivated further analyses of parasitised DCs under shear stress.

**Figure 3.**
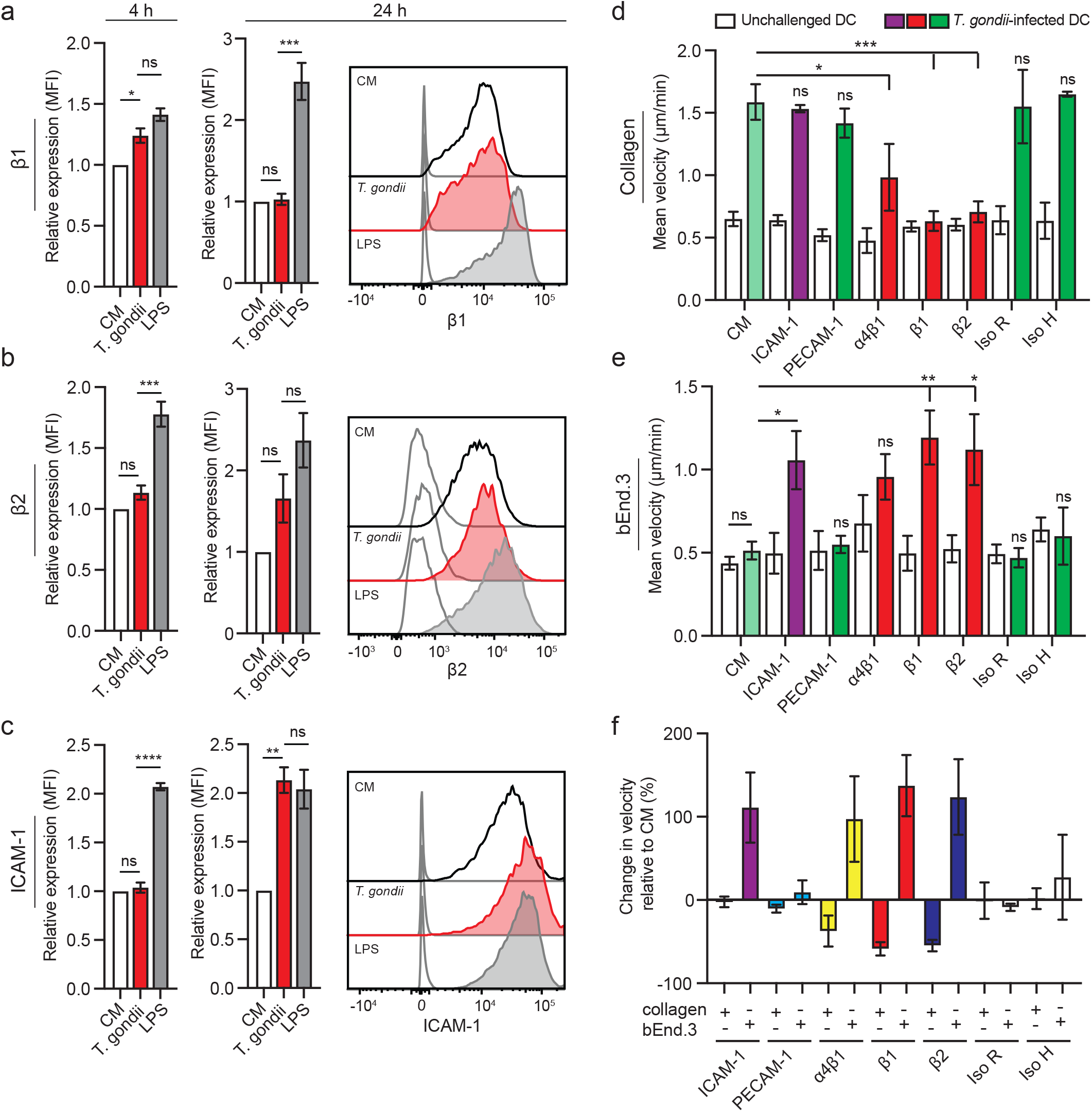
Motility of *T. gondii*-infected DCs upon blockade of integrins and CAMs. **(a, b, c)** Bar graphs show the relative expression of β1 (CD29), β2 (CD18) integrins and ICAM-1 (CD54), respectively, at 4 and 24 h post-challenge of CD11c^+^ cells with *T. gondii* tachyzoites (GFP-expressing ME49, MOI 1) or LPS (100 ng/ml), assessed by flow cytometry (see Supplementary Fig. S2 for gating strategy). For each time point, the mean fluorescence intensity (MFI) was related to that of unchallenged CD11c^+^ cells in complete medium (CM, normalised to 1). Histograms show fluorescence intensity distributions for indicated cell populations/conditions and for corresponding isotype controls (unfilled curve with grey line). **(d, e)** Mean velocity of unchallenged and *T. gondii*-challenged DCs treated with inhibitors and antibodies as stated, on (a) collagen or (b) bEnd.3 monolayers, respectively. CM indicates complete medium. **(f)** Percentage (%) change in velocity of *T. gondii*-challenged DCs compared to unchallenged DCs for each condition. All data are presented as mean ± s.e.m from 3 independent experiments (n = 3) \**P* < 0.05, \*\**P* < 0.01, \*\*\**P* < 0.001, \*\*\*\**P* < 0.0001, ns; non-significant by one-way ANOVA, Dunnett’s post-hoc test.

### Adhesion and motility of *T. gondii*-challenged DCs under flow conditions

To assess DC motility under more physiological conditions, unchallenged or *T. gondii*-challenged DCs were tracked under shear stress in flow chambers coated with collagen or bEnd.3 cells (**Fig. 4a, b**). On collagen, the numbers of adhered DCs were reduced upon *T. gondii* challenge (**Fig. 4c**). The mean pathlength of *T. gondii*-challenged DCs (related to unchallenged) was increased, and their mean velocities were elevated following 0.2 dyn/cm^2^ shear stress conditions (**Fig. 4d, e**). However, non-significant differences were observed between unchallenged and *T. gondii*-challenged DCs at 1 dyn/cm^2^, indicating detachment at higher shear stress (**Fig. 4d, e**). In contrast, on bEnd.3 cell monolayers, adhesion of *T. gondii*-challenged DCs was elevated compared with unchallenged DCs following 0.2 dyn/cm^2^ flow, with non-significant differences after 1 dyn/cm^2^ (**Fig. 4f**). While pathlengths and velocities remained high compared to collagen condition, non-significant differences in pathlength or velocity were observed between unchallenged and *T. gondii*-challenged DCs (**Fig. 4g, h**), in line with observations under static conditions (Fig 2c). Thus, under shear stress, collagen and endothelium had opposite effects on the adhesion of parasitised DCs by reducing and elevating numbers of adhered cells, respectively, related to unchallenged DCs (**Fig. 4i**). Moreover, the pathlengths of adherent parasitised DCs were elevated in collagen but not on endothelium at low shear stress, related to unchallenged DCs (**Fig. 4j**). Altogether, this extends and corroborates results under static conditions (Figs. 2 and 3). Jointly, the data indicate that the *T. gondii*-induced hypermotility of DCs is maintained on collagen under low shear stress but is abolished upon binding of parasitised DCs to the endothelium. Nonetheless, adherent parasitised DCs were motile on endothelial monolayers.

**Figure 4.**
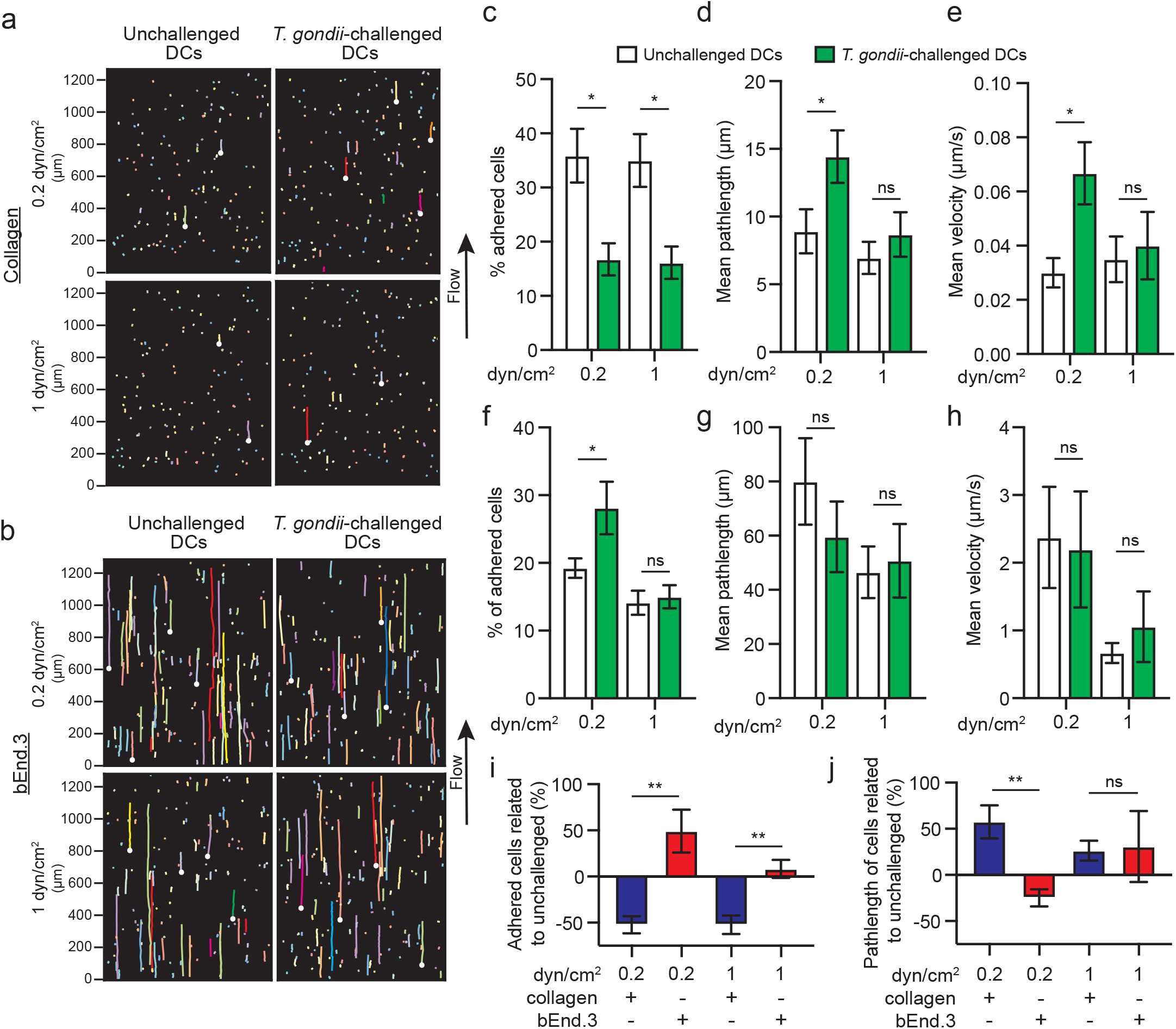
Adhesion and motility of *T. gondii*-challenged DCs under flow conditions. (**a, b**) Representative cell traces of unchallenged or *T. gondii*-challenged DCs on (a) collagen or (b) bEnd.3 cell monolayers, at indicated shear stress (dyn/cm^2^). Each color-coded trace shows the euclidian displacement distance of a single cell. White dots indicate the start point of randomly selected track lengths, showing that cells migrated in the direction of flow. **(c)** Percentage (%) of DCs adhered to collagen, following fluidic shear stress (dyn/cm^2^), related to total number of DCs in the same frame in static condition as indicated under Materials and Methods. **(d, e)** Mean pathlength and velocity, respectively, of tracked cells on collagen. **(f)** Percentage (%) of DCs adhered to bEnd.3 cell monolayers, following fluidic shear stress (dyn/cm^2^), related to total number of DCs in the same frame in static condition. **(g, h)** Mean pathlength and velocity, respectively of tracked cells on bEnd.3 cell monolayers. **(i, j)** Percentage change in adherence and velocity, respectively of *T. gondii*-challenged DCs related to unchallenged DCs, for each condition. Bar graphs show mean ± s.e.m. from 4 independent experiments (n = 4). * *P* < 0.05, ** *P* < 0.01; ns, non-significant by one-way ANOVA, Sidaks post-hoc test (c, d, e, f, g, h) or Students’ *t*-test (i, j).

### Implication of CAMs and integrins in the transmigration of *T. gondii*-challenged DCs across endothelium

Because CAMs and integrins play a central role in the extravasation of leukocytes across endothelium [13], we assessed the effects of CAM and integrin blockade on the TEM of unchallenged and *T. gondii*-challenged DCs across polarised bEnd.3 cell monolayers. First, we performed a transcriptional profiling for relevant integrins and CAMs (**Supplementary Fig. S3**) and controlled that the treatments non-significantly impacted permeability (**Fig. 5a, b**) and TEER (**Fig. 5c, d**). Interestingly, blockade of ICAM-1, PECAM-1, α4β1, β1 and β2 integrins significantly reduced the TEM frequency of *T. gondii*-challenged DCs (**Fig. 5e**), but non-significantly impacted the TEM frequency of unchallenged DCs (**Fig. 5f**). Jointly with motility assays (Fig. 3), this indicates that parasitised DCs utilise integrin and CAM interactions to adhere and perform TEM, which is potentiated by *T. gondii* infection.

**Figure 5.**
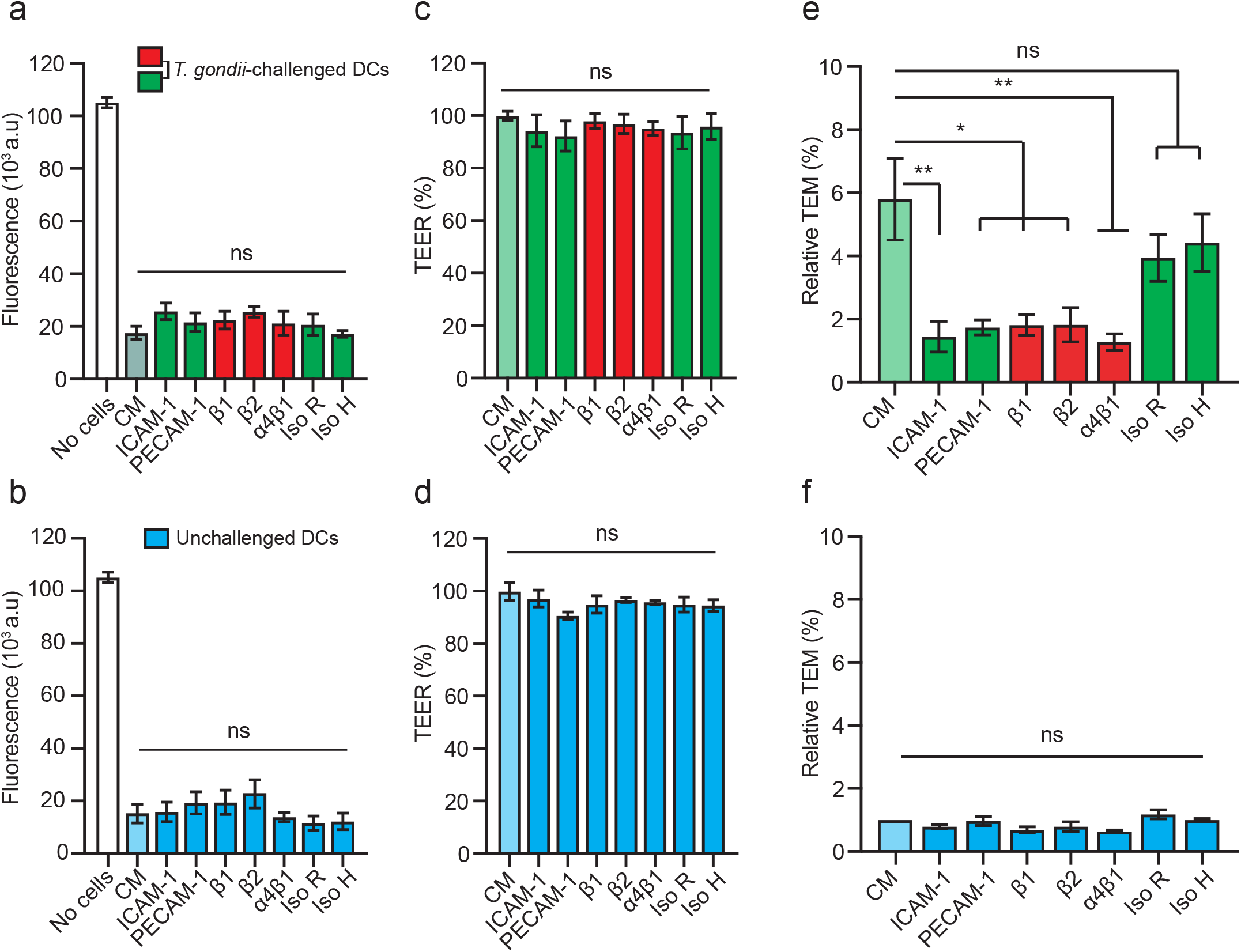
TEM of *T. gondii*-challenged DC upon integrin CAM blockade. **(a, b)** Permeability of bEnd.3 monolayers to FITC-dextran (3 kDa) after 16 h exposure to *T. gondii*-challenged DCs or unchallenged DCs ± antibodies or inhibitors, respectively, and as indicated. CM indicates complete medium. **(c, d)** TEER (%) of bEnd.3 monolayers as in (a) and (b), respectively. **(e, f)** Relative TEM (%) of *T. gondii*-challenged DCs or unchallenged DCs, respectively. Relative TEM frequency of DCs across bEnd.3 monolayers are shown as percentage (%) of DCs added, normalised to unchallenged DC condition for each experimental condition (set to 1.0, light blue bar ‘CM’). Data are presented as mean ± s.e.m. from 4 independent experiments (n = 4). \**P* < 0.05, \*\**P* < 0.01, ns; non-significant by one-way ANOVA, Dunnett’s post-hoc test (a-f).

### Gene silencing of talin-1 (*Tln1*) or of β1 integrin (*Itgb1*) impacts motility and TEM of *T. gondii*-infected DCs on endothelium under static and flow conditions

Consistent with paradigms of DC migration [5], we previously showed that amoeboid migration of parasitised DCs in collagen matrix is an integrin-independent process [26]. However, extravasation of DCs across endothelium is integrin-dependent [16] and integrins contribute to the migration of parasitised DCs [30]. To further assess the role of integrins in motility and TEM of *T. gondii*-infected primary DCs, we gene silenced the integrin-linking protein talin (Tln1) and β1 integrin (ITGB1). In transduced DCs, significant reductions in *Tln1* and *Itgb1* mRNA expression, respectively, were measured, related to control shLuc-transduced DCs (**Fig. 6a, b**). First, to confirm the impact of gene silencing on integrin-mediated motility, transduced cells were challenged with *T. gondii* and allowed to migrate on collagen-coated surfaces (**Supplementary Fig. S4**). Motility and mean velocities of shITGB1 and Tln1-transduced *T. gondii*-infected DCs were significantly reduced on collagen (**Fig. 6c, d**). In contrast, on bEnd.3 cell monolayers, shITGB1- and Tln1-transduced *T. gondii*-infected DCs exhibited increased velocities compared to mock and shLuc-transduced cells (**Fig. 6e, f**). A similar effect was confirmed under flow, with reduced adhesion (Fig. 6g) and elevations in pathlengths and velocities of transduced infected DCs (Fig. 6h, i). Importantly, the TEM frequency of *T. gondii*-challenged DCs across polarised endothelial monolayers **(Fig. 6 j, k)** was significantly decreased in shITGB1 and shTln1-transduced cells compared to shLuc- or mock-transduced cells (**Fig. 6l**). We conclude that gene silencing of talin-1 (*Tln1*) or of β1 integrin (*Itgb1*) inhibits hypermotility and TEM but elevates the motility of *T. gondii*-infected DCs on endothelium.

**Figure 6.**
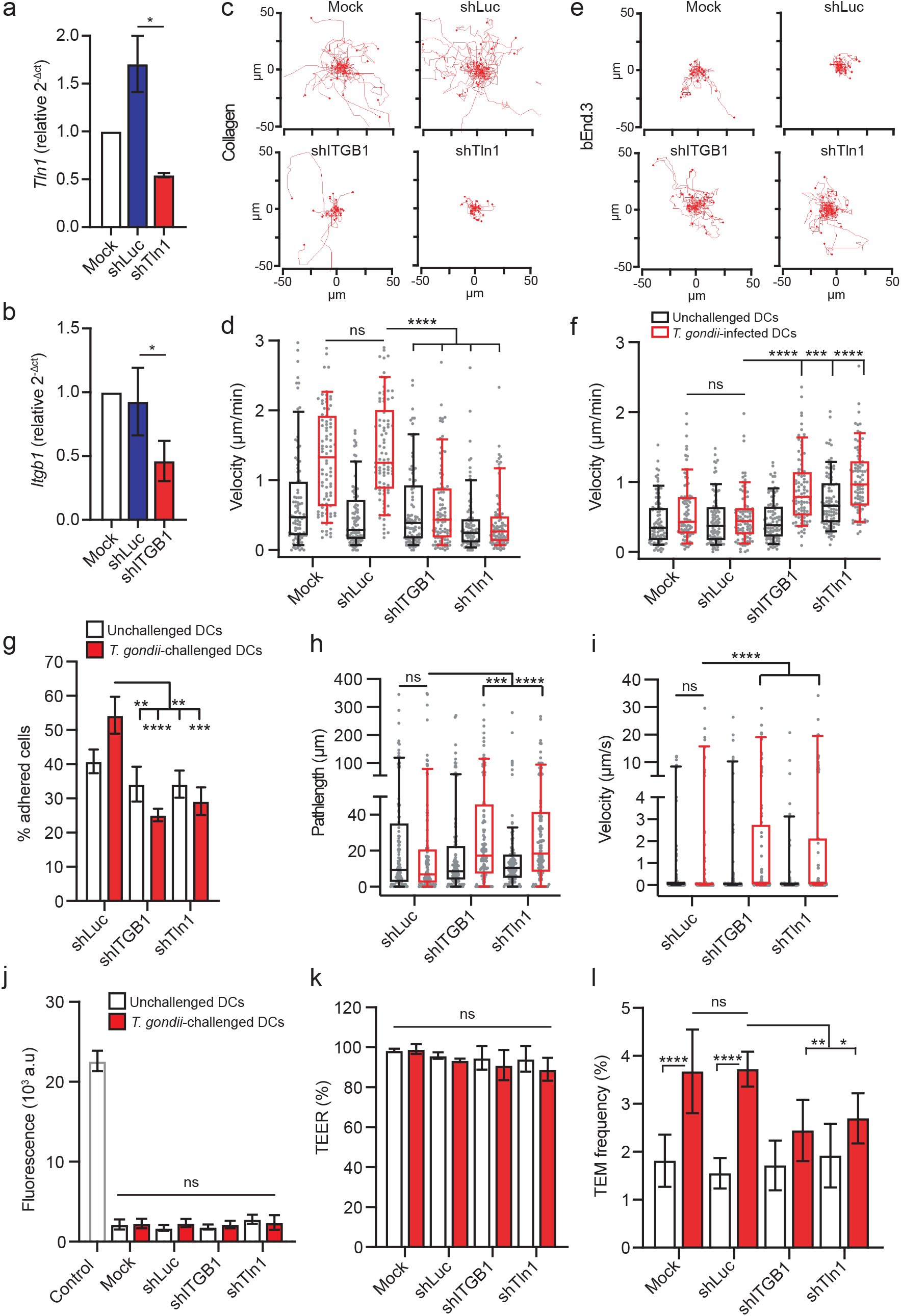
Impact of gene silencing of talin (*Tln-1*) and β1 integrin (*Itgb1*) on the motility and transmigration of *T. gondii*-challenged DCs. **(a)** *Tln1* mRNA expression (2^-ΔCt^) in DCs transduced with shTln1 or control shLuc lentivirus relative to mock-treated cells (mock set to 1.0). **(b)** *Itgb1* mRNA expression (2^-ΔCt^) in DCs transduced with shITGB1 or control shLuc lentivirus relative to mock-treated cells. **(c)** Representative motility plots of mock-treated DCs and DCs transduced with lentiviral vectors targeting *Itgb1* mRNA (shITGB1), *Tln1* mRNA (shTln1) or a non-expressed target (shLuc) challenged with *T. gondii*, on collagen. **(d)** Box-and-whisker and scattered dot plots represent median velocities (μm/min) of DCs on collagen, as in (c). Grey circles represent velocities from individual cells. **(e)** Representative motility plots of mock-treated DCs and DCs transduced with lentiviral vectors targeting *Itgb1* mRNA (shITGB1), *Tln1* mRNA (shTln1) or a non-expressed target (shLuc) challenged with *T. gondii*, on bEnd.3 monolayers. **(f)** Box-and-whisker and scattered dot plots represent median velocities (μm/min) of DCs on bEnd.3, as in (e). Grey circles represent velocities from individual cells. **(g)** Bar graphs show percentage (%) of transduced DCs adhered to bEnd.3 monolayers related to DCs in the same frame in static condition, following fluidic shear stress (0.2 dyn/cm^2^). **(h, i)** Box-and-whisker and scattered dot plots represent median pathlengths (μm) and velocities (μm/s), respectively, of tracked cells on bEnd.3 monolayers. Grey circles represent pathlengths or velocities, respectively, from individual cells. **(j, k)** Permeability to FITC-dextran (3 kDa) and TEER (%), respectively, of bEnd.3 cell monolayers following DC transmigration. **(l)** TEM frequency shown as percentage (%) of DCs added in the upper well. Bar graphs show mean ± s.e.m and box plots show median from 3-4 independent experiments (n = 3-4). In box-and-whisker plots, boxes mark 25^th^ to 75^th^ percentile and whiskers mark 10^th^ and 90^th^ percentiles of the datasets. * *P* < 0.05, \*\**P* < 0.01, \*\*\**P* < 0.001, **** *P* < 0.0001; ns, non-significant by Students’ *t*-test (a, b), One-way ANOVA, Dunnett’s post-hoc test (d, f), Kruskal-Wallis, Dunn’s post-hoc test (h, i), repeated measures one-way ANOVA, Sidak’s post-hoc test (j, k, l).

## Discussion

The interconversion between migratory states of leukocytes and the crosstalk between CAMs and integrins are crucial for diapedesis [16]. Here, we explored the interplay between endothelium and *T. gondii*-challenged DCs and found that intracellular parasitisation impacts migration in a cell environment-related fashion, with a pivotal role for integrins.

We report that infection with *T. gondii* induces dramatic migratory changes in primary DCs, which impact on locomotion, adhesion and enhanced transmigration across polarised endothelial cell monolayers. Upon *T. gondii* infection, DCs undergo MAT in extracellular matrix and enhanced amoeboid migration -termed hypermotility- [26,24,30,31], which promotes parasite dissemination *in vivo* [22,28]. Surprisingly, we found that the hypermotility of parasitised DCs in 3D and 2D collagen confinements (**Fig 7a, b**) is lost upon direct contact with endothelium (**Fig. 7c**), and that the mean velocities and migrated distances became comparable between unchallenged and *T. gondii*-infected DCs. The reduction of motility of DCs on endothelium suggests instead, arrest and adhesion, which is regulated by integrin-CAM crosstalk [37,16], and precedes leukocyte TEM (**Fig 7d**).

**Figure 7.**
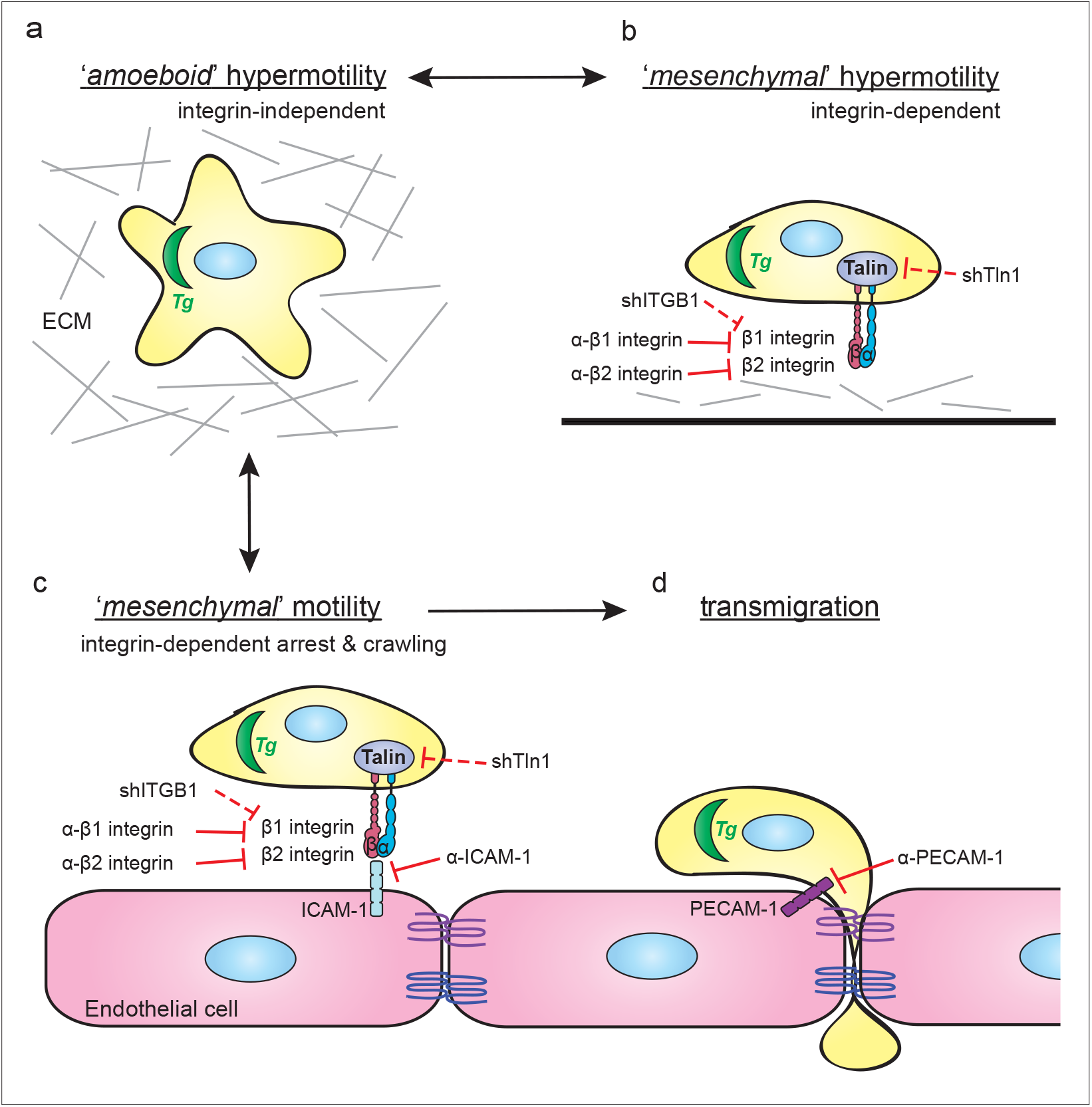
Schematic representation of the integrin-dependent motility switches in *T. gondii*-infected DCs leading to transendothelial migration. **(a)** *T. gondii* (*Tg*) actively invades DCs and induces rapid integrin-independent DC migration, termed hypermotility, in 3-dimensional (3D) extracellular matrix (ECM) confinements, such as collagen. *T. gondii*-induced hypermotility has features of amoeboid motility [26]. **(b)** The hypermotility of *T. gondii*-infected DCs on 2D collagen-coated surfaces has the adhesion/integrin-dependency feature of mesenchymal migration, however with abrogated proteolysis [24,30]. Blockade of β1 or β2 integrin, or gene silencing of talin (shTln1) or β1 integrin (shITGB1) abolishes hypermotility of *T. gondii*-infected DCs on collagen. **(c)** On endothelium, *T. gondii*-infected DCs adhere to cell monolayers under static and flow conditions. In relation to hypermotility on collagen and in ECM (a, b), the reduced motility by parasitised DCs on endothelium is integrin- and CAM-dependent, with features of mesenchymal migration. Specifically, Ab blockade of β1 or β2 integrins, or gene silencing of talin (shTln1) or β1 integrin (shITGB1) elevates *T. gondii*-infected DC motility on endothelium, while the same treatments reduce motility on collagen-coated surfaces (b). **(d)** *T. gondii*-infected DCs exhibit elevated transmigration across polarised endothelial monolayers, related to non-infected DCs or infected monocytes/macrophages. Ab blockade of β1 or β2 integrins, ICAM-1, PECAM-1 or gene silencing of talin (shTln1) or β1 integrin (shITGB1) decreases the transmigration frequency of *T. gondii*-infected DCs across endothelium.

Under shear stress conditions, parasitised DCs elevated or maintained adhesion to endothelium, while adhesion was conversely reduced on collagen. The elevated adhesion on endothelium is in contrast with the MAT-related morphological changes that parasitised DCs present on collagen and in collagen matrix, including round-cell morphology, decreased adhesion and redistribution of integrins [25,26,24]. Moreover, under flow conditions, the pathlengths of parasitised DCs were maintained high with reduced velocities, indicating crawling of DCs [16]. *In vivo*, leukocyte diapedesis mainly occurs at the postcapillary venules of inflamed tissue, at low shear stress levels [13]. Further, exposure to shear stress can increase transcription levels of ICAM-1, PECAM-1 and other endothelial CAMs [12], which interact with leukocyte integrins for diapedesis. This motivated a further investigation of the role of integrins.

We demonstrate that integrin signalling plays a pivotal role in the infection-related arrested motility of DCs on endothelium. Interestingly, disruption of the integrin-CAM interaction by blockade of β1 and β2 integrins, along with ICAM-1, partially restored the hypermotility of infected DCs on endothelium. In contrast, on collagen-coated surfaces, the hypermotility phenotype in *T. gondii*-infected DCs was reduced upon β1 and β2 integrin blockade, and gene silencing of β1 integrin or talin. We recently reported a role for β1 integrin in a motogenic TIMP1-CD63-ITGB1-FAK signalling axis associated with MAT [30]. Here, we add that β1 integrin is equally important for, presumably, amoeboid-to-mesenchymal migratory conversion and adhesion to endothelium. Not in contraposition with the above, amoeboid migration of DCs is integrin-independent in ECM [5], and integrin-blocking antibodies non-significantly impacted the migration of *T. gondii*-infected DCs in a 3D collagen matrix [26]. More recent studies have also shown that talin-deficient T cells, lacking integrin-based force transmission, are unable to adhere to or migrate on 2D surfaces [38]. Similarly, the switch of cancer cells between amoeboid and mesenchymal migration modes appears to be advantageous in tumour invasion, depending on their extracellular environment [39]. Migrating leukocytes have the ability to slow down or speed up migration in seconds, switching from high-speed actin-flow driven motility, to slow adhesion-driven motility mediated by integrins [4]. The difference in motility observed between parasitised DCs on collagen versus endothelium, implies a pivotal role for integrins in each scenario. It also reflects the ability of DCs to switch between different modes of migration depending on the extracellular environment (**Fig. 7**). On the one hand, infection-induced hypermotility in matrix is linked to increased dissemination [28,29,23], but upon reaching barriers such as the BBB, adhesion is necessary for TEM and thus facilitates parasite dissemination across the vasculature.

Mesenchymal cell migration is characterized by cell polarisation, adhesive interactions with the substratum and proteolytic extracellular matrix remodelling, followed by retraction of the cell rear to achieve cellular movement [6]. In fact, the motility of parasitised DCs on 2D collagen or endothelial cell surfaces fulfils few criteria to be classified as strictly mesenchymal. Our data show that this motility relies on adhesive interactions, in contrast to motility in 3D matrix which is integrin-independent [26]. However, a number of observations also argue against strict mesenchymal motility. Specifically, parasitised DCs abrogate pericellular matrix proteolysis by secretion of TIMP-1 [24] and undergo gross morphological changes, encompassing rounding-up and irreversible dissolution of podosomes with redistribution of integrins that are, in fact, more consistent with amoeboid motility [25]. The intracellular parasite achieves migratory activation of DCs by interfering with MAP kinase signalling [30,31] and calcium signalling [29] via activation of GABAergic signalling in the infected cell [28,23]. Here, we add that the hypermigration of parasitised DCs [40] is modulated by external cues in the cellular environment. In the blood circulation, it is unlikely that the sequestration of parasitised DCs to endothelium solely depends on the migratory mode *per se* but rather on the ability to adhere. Thus, that hypermotile parasitised DCs adhere to brain endothelial monolayers in an integrin-dependent fashion bears relevance for concepts of parasite dissemination.

We demonstrate that integrins contribute to the enhanced TEM of *T. gondii*-infected DCs across polarised primary brain endothelial cell monolayers (**Fig. 7d**). Interestingly, transmigration occurred without measurable perturbation of polarisation or disruption of the cell barrier integrity, in line with previous reports [41] and indicative of regulated cell-cell interactions. Indeed, the TEM of infected DCs was reduced upon blockade of adhesion molecules, ICAM-1 and PECAM-1, and β1 and β2 integrins. Moreover, gene silencing of the integrin-cytoskeleton linker talin in DCs demonstrated a crucial role for integrins. Taken together with the motogenic role of β1 integrin [30], here we add a role in adhesion to endothelium and TEM. Thus, while β2 integrins contribute, we identify a pivotal role for β1 integrins. Jointly, this shows that, potentiated by *T. gondii* infection, integrin-CAM interactions mediated TEM. It remains largely unknown how intracellular pathogens modulate the migration modes of shuttle-leukocytes to enter the CNS, especially initially during infection in the absence of CNS inflammation [42]. However, integrin-CAM interactions are likely to be in play, as shown here for *T. gondii*-infected DCs. A recent report using a fungal infection model in mice showed that α4β1 integrin (VLA-4) mediated recruitment of infected monocytes to the BBB, which subsequently migrated to the brain parenchyma [43]. In HIV infection, disrupting the interaction between αLβ2 (LFA-1) on CD4^+^ T cells and ICAM-1 on DCs by mAbs inhibited viral transmission between T cells and DCs [44,45], and thus similar interactions may be implicated at the BBB. Beyond infection, the pathogenesis of multiple sclerosis implicates a recruitment of DCs and T cells to the CNS [46] and has opened up for mAb-based therapies targeting integrins, for example α4β1 [47]. Of note, parasitised DCs, but not by-stander DCs, elevated their expression of ICAM-1, indicating effects related to intracellular parasitation. Further, this also indicates that blockade of ICAM-1 in our study may inhibit adhesion mediated by both DC and endothelial cell ICAM-1 [48,49]. Reciprocally, blockade of both DC and endothelial integrins may also take place [50-52]. Additionally, future studies need to address if the infection-induced cytoskeletal changes with redistribution of integrins alter adhesion [25] and if the activation state of integrins is altered in parasitised DCs [53]. Also, given the profound impact of talin on the migratory responses described here, its role in integrin activation and mechanotransduction in parasitised DCs awaits further investigation [15].

Our data shows that the TEM frequency of parasitised leukocytes not only depends on the cellular environment but also is related to cell type. The TEM frequency of *T. gondii*-infected macrophages, derived from the same donor mice, was 10-20-fold inferior to that of infected DCs. For human monocytes, comparable transmigration frequencies have been reported for unchallenged and *T. gondii*-challenged monocytes across filters and non-polarizing endothelial monolayers [54,53]. *T. gondii*-challenged monocytes have also been reported to exhibit hypermotility but reduced transmigration [55], contrasting with the elevated transmigration of parasitised DCs and microglia [22,56]. Further, elevated TEM was observed in infected rat peripheral blood mononuclear cells (PBMCs) [57] and, under shear stress conditions on HUVEC monolayers, the motility and TEM frequencies of infected monocytes increased compared to static conditions [53,58]. Altogether, this highlights that the cellular environment likely impacts migratory responses in a cell type-related fashion [59] and that DCs may provide dissemination advantages over other mononuclear phagocytes.

In experimental mouse toxoplasmosis, the transportation of *T. gondii* by DCs and other monocytic cells in the circulation has been associated with higher parasitic loads in the CNS [21,22]. DCs migrate into the brain parenchyma in an integrin-dependent fashion during toxoplasmosis [60]. However, it remains unclear if DCs can transport *T. gondii* into the CNS, as extracellular parasites can also invade and transmigrate across endothelium [61,36]. Here, we provide *in vitro* proof-of-concept that transmigration of DCs across highly-polarised primary endothelium is potentiated by infection. In a broader context, leukocytes can be activated to perform diapedesis by chemotactic signalling [62]. However, activation does not automatically imply enhanced TEM. For example, upon LPS-stimulation, both reduced and elevated TEM by leukocytes have been reported [63-65]. The role of chemokines in homing of leukocytes to the CNS remains partly unelucidated [66]. Also, natural primary *T. gondii* infection of humans and vertebrates is asymptomatic or accompanied by mild symptomatology. Thus, the initial penetration of *T. gondii* to the CNS likely happens in absence of generalized inflammation or reactive leukocyte infiltration to the parenchyma [67]. Nevertheless, it remains to be investigated if inflammatory mediators, such as IFN-γ and IL-12, impact the interaction of endothelium with parasitised leukocytes. *T. gondii* induces amoeboid migration of DCs and other mononuclear phagocytes by activating alternative signalling pathways, including GABAergic signalling and MAP kinase activation [23,31], which can synergise with chemotaxis in vitro [28,25]. Thus, we postulate that the infection-related enhanced TEM shown here *in vitro* may facilitate the diapedesis of parasitised DCs to the brain parenchyma, synergistically with chemotactic and inflammatory cues. The elucidation of alternative activation pathways that can drive leukocyte TEM will further our knowledge on how leukocytes and other migratory cells regulate their migratory states for vital biological processes.

## Declarations

### Ethics approval

The Regional Animal Research Ethical Board, Stockholm, Sweden, approved protocols involving extraction of cells from mice, following proceedings described in EU legislation (Council Directive 2010/63/EU).

### Data availability

The datasets used and/or analysed in this study are available from the corresponding author on reasonable request.

### Author’s contributions

E.C.R and ALH performed experiments and analysed the data. All authors contributed to the generation of figures and writing of this manuscript.

### Code availability

not applicable

## Acknowledgements

We thank members of the Barragan lab, Manuel Varas-Godoy, University of Santiago, Chile for critical input and Sascha Granberg for graph generation in Python.

## Supplementary material

**Supplementary Table S1. Sequences for shRNAs**.

**Supplementary Table S2. Sequences for qPCR primers**.

**Supplementary Figure S1. Immunostainings of endothelial tight junctions**

**(a, b**) Representative micrographs of primary mouse brain endothelial cells (MBECs) stained for (**a**) ZO-1 (red), (**b**) occludin (red), with DAPI-stained nuclei (blue). Scale bars = 50 μm.

**Supplementary Figure S2. Gating strategy for flow cytometry and characterizations of CD11c**^**+/-**^ **cells and by-stander cells**.

**(a)** Bivariate dot plots show gating strategy for the experiments displayed in Figure 3, exemplified with bone marrow-derived cells challenged with GFP-expressing *T. gondii*. Histogram shows fluorescence intensity distributions and gates used to define CD11c^+^ cells. Numbers indicate percentage (%) of cells in corresponding gate.

**(b)** Bivariate contour plots of live unchallenged DCs stained for CD11c vs. β1, β2 integrins or ICAM-1, respectively.

**(c, d)** Bar graphs show the relative expression of β1 (CD29), β2 (CD18) integrins and ICAM-1 (CD54), respectively, at 4 h (c) and 24 h (d) post-challenge of cells (CD11c^+^ and CD11c^-^) with *T. gondii* tachyzoites (GFP-expressing ME49, MOI 1) or LPS (100 ng/ml), assessed by flow cytometry. For each time point and condition, the mean fluorescence intensity (MFI) was related to unchallenged cells in complete medium (CM, normalised to 1). ‘By-stander’ indicates GFP^-^ cells (non-infected) in cell suspensions challenged with *T. gondii*.

Data are presented as mean ± s.e.m from 3 independent experiments (n = 3) \*\**P* < 0.01, \*\*\*\**P* < 0.0001, ns; non-significant by two-way ANOVA, Sidak’s post-hoc test.

**Supplementary Figure S3. Transcriptional analyses of bEnd.3 cells and DCs**

**(a)** mRNA expression of indicated genes in bEnd.3 cells challenged with *T. gondii*, unchallenged DCs, *T. gondii*-challenged DCs or LPS, relative to CM condition, after 6 h.

**(b)** mRNA expression of unchallenged bEnd.3 cells and DCs.

Bar graphs show mean ± s.e.m. from 2-3 independent experiments. * *P* < 0.05, \*\**P* < 0.01, \*\*\**P* < 0.001, \*\*\*\**P* < 0.0001; ns, non-significant by two-way ANOVA, Dunnett’s post-hoc test.

**Supplementary Figure S4. Transductions of primary DCs with shRNAs** Transductions were performed as indicated under Materials and Methods. **(a-d)** For each condition, representative micrographs show unchallenged DCs (left) and *T. gondii*-challenged DCs (right), mock-treated or treated with control shRNA (shLuc) or shRNA to indicated target gene. Transduced cells were distinguished by expression of the reporter eGFP (green) and *T. gondii* by expression of RFP (red). Infected cells were defined as green + red cells (indicated by white arrows). Scale bar = 100 μm.

