## Supplementary material for "Integrin-dependent migratory switches regulate the translocation of *Toxoplasma*-infected dendritic cells across brain endothelial monolayers": Supplenat Tables 1 and 2, Suppl. Figs. S1-4

#### Table S1. Sequences for shRNAs.

| Target | Sequence (5' to 3') |
|---|---|
| shLuc: | TGTTCTCCGAACGTGTCACGTTTCAAGAGAACGTGACACGTTCGGAGAACTTTTTTC |
| shITGB1: | CCGGGCCATTACTATGATTATCCTTCTCGAGAAGGATAATCATAGTAATGGCTTTTTG |
| shTln1: | TGCTGGGAAAGCTTTGGACTACTACTTCAAGAGAGTAGTAGTCCAAAGCTTTCCCAGCTTTTTTC |

#### Table S2. Sequences for qPCR primers.

| Target | Primer pair sequence (5' to 3') |
|---|---|
| *Tln1* | Fd: GGTGAAGACTATCATGGTGG<br>Rv: TTGGTGATACCAATTCGGG |
| *Itgb1* | Fd: GATGAATTTGCAACTGGTTTCC<br>Rv: GCAAGATTTGGCATTTGCT |
| *Gapdh* | Fd: TGACCTCAACTACATGGTCTACA<br>Rv: CTTCCCATTCTCGGCCTTG |
| *Vcam1* | Fd: GTGACTCCATGGCCCTCACTT<br>Rv: CGTCCTCACCTTCGCGTTTA |
| *Icam1* | Fd: CAATTTCTCATGCCGCACAG<br>Rv: CTGGAAGATCGAAAGTCCGG |
| *Sele (E-selectin)* | Fd: CCCTGCCCACGGTATCAG<br>Rv: ACGTGCATGTCGTGTTCCA |
| *Itga4* | Fd: ATGGCTGCGGAAGCGATGTGC<br>Rv: CATGCCATAGCAAACACCAGTGG |
| *Itgb2* | Fd: CAGGAATGCACCAAGTACAAAGT<br>Rv: CCTGGTCCAGTGAAGTTCAGC |
| *CD31* | Fd: CTGCCAGTCCGAAAATGGAAC<br>Rv: CTTCATCCACCGGGGCTATC |
| *Cldn5* | Fd: TTTCTTCTATGCGCAGTTGG<br>Rv: GCAGTTTGGTGCCTACTTCA |
| *Itgax* | Fd: CTTCCCAGACTTGAAGACC<br>Rv: TCTTCTCCATCATTAGACACC |

### Figure S1

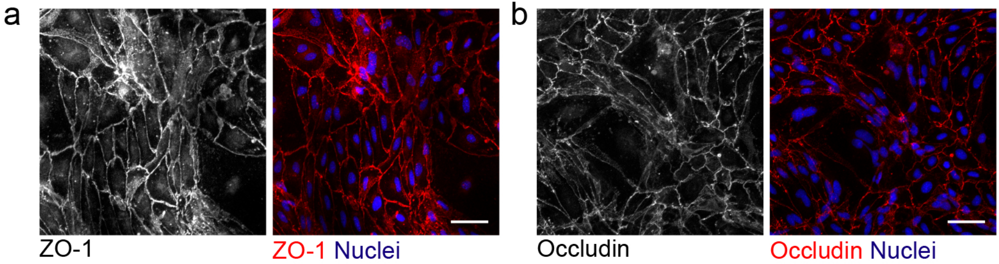

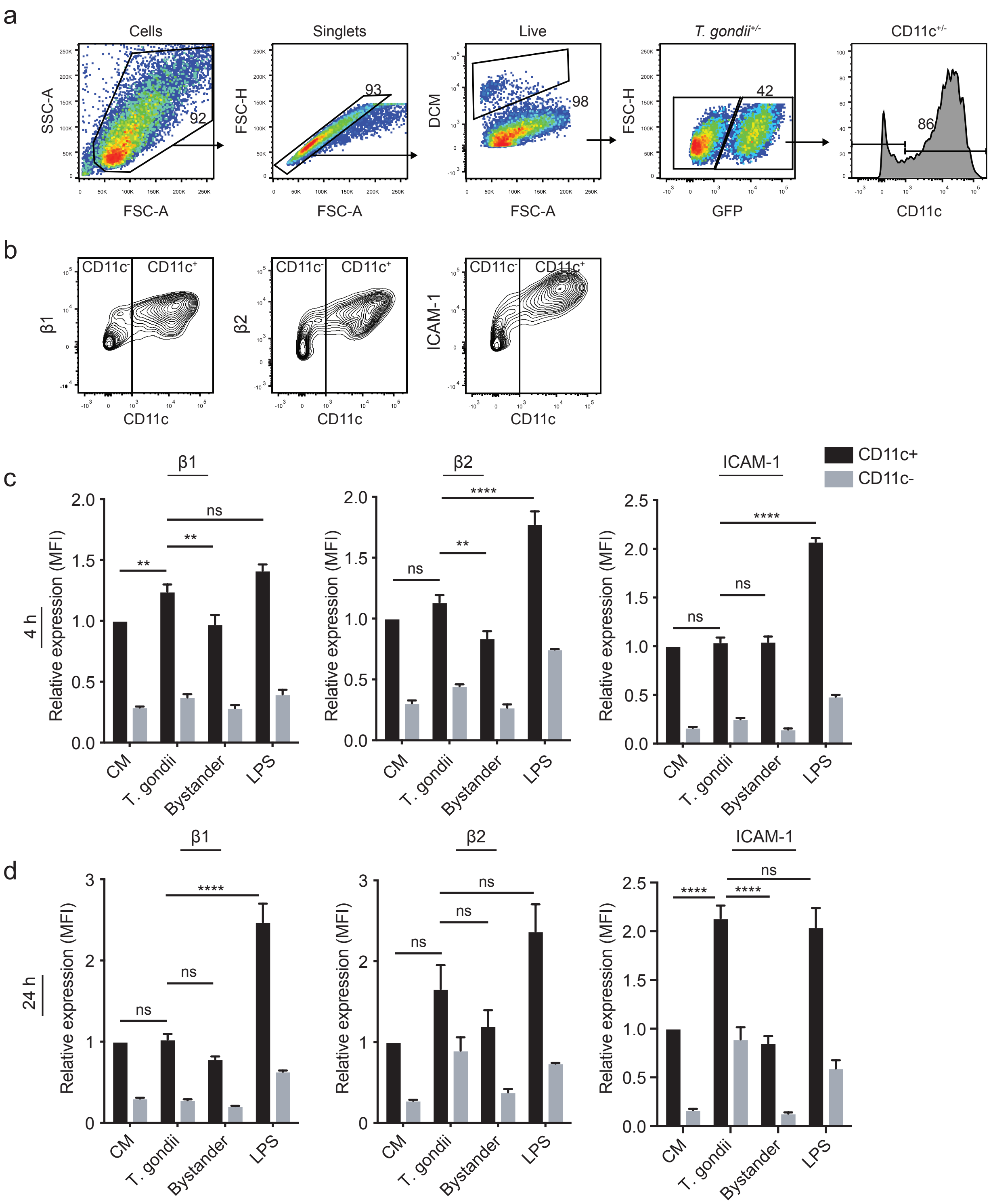
a
Cells
Singlets
Live
T. gondii+/-
CD11c+/-
SSC-A
FSC-A
FSC-H
DCM
GFP
CD11c
92
93
98
42
86
b
CD11c-
CD11c+
β1
β2
ICAM-1
c
β1
β2
ICAM-1
CD11c+
CD11c-
4 h
Relative expression (MFI)
CM
T. gondii
Bystander
LPS
ns
**
****
d
24 h
ns
****

### Figure S3

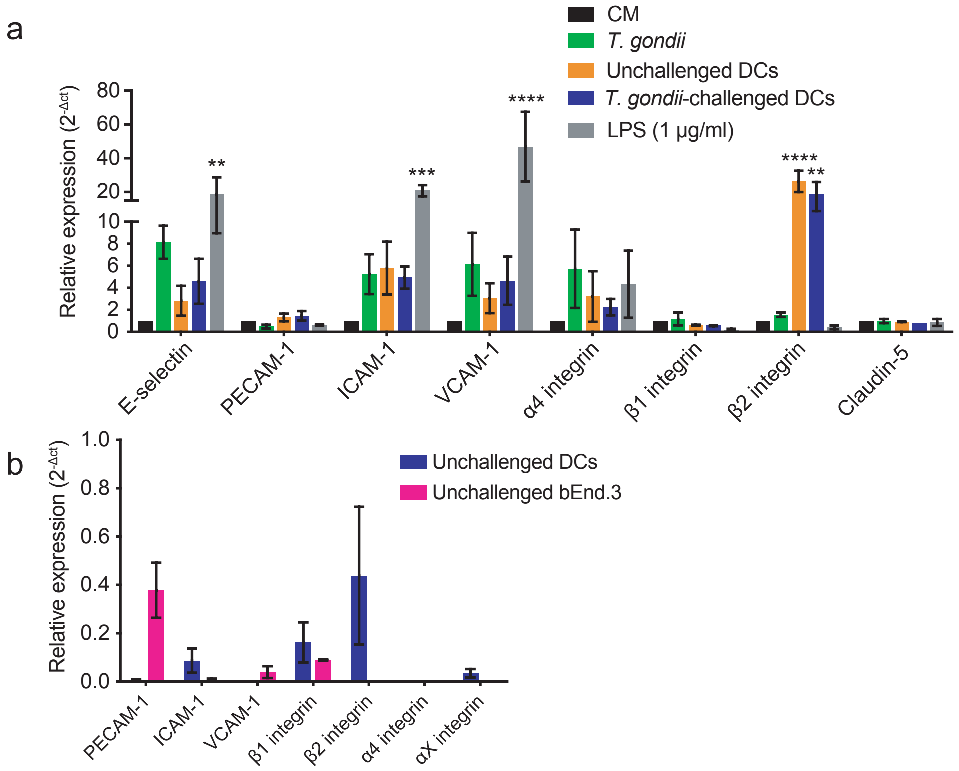
a
CM
T. gondii
Unchallenged DCs
T. gondii-challenged DCs
LPS (1 µg/ml)
Relative expression (2-Δct)
80
60
40
20
10
8
6
4
2
0
**
***
****
****
**
E-selectin
PECAM-1
ICAM-1
VCAM-1
α4 integrin
β1 integrin
β2 integrin
Claudin-5
b
Unchallenged DCs
Unchallenged bEnd.3
Relative expression (2-Δct)
1.0
0.8
0.6
0.4
0.2
0.0
PECAM-1
ICAM-1
VCAM-1
β1 integrin
β2 integrin
α4 integrin
αX integrin

Figure S4

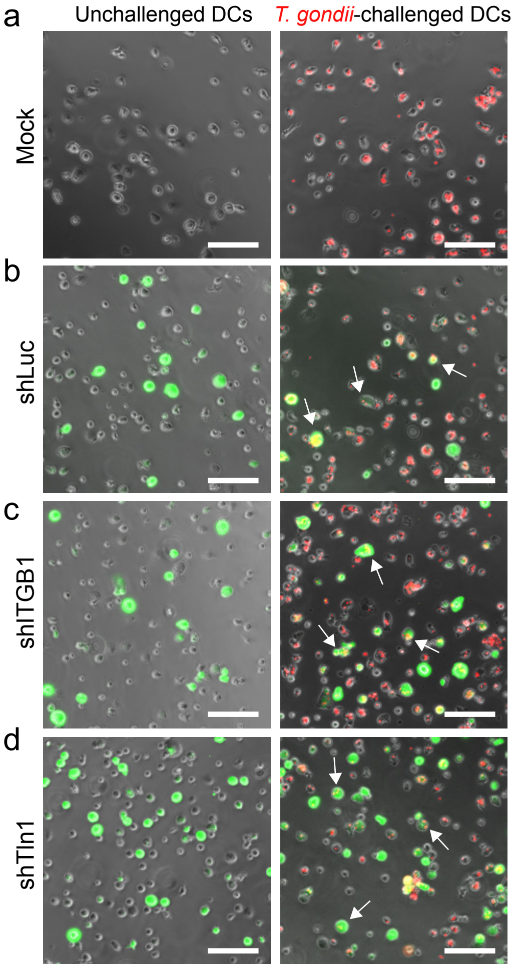
a
Unchallenged DCs
T. gondii-challenged DCs
Mock
b
shLuc
c
shITGB1
d
shTln1
